# IL-12/IL23 blockade reveals patterns of asynchronous inflammation in pyoderma gangrenosum

**DOI:** 10.1101/2024.04.26.591387

**Authors:** Rashi Yadav, Morgan Vague, Malia Rettig, Christopher P. Loo, Kasidy Brown, Abrar Samiea, Joshua M. Moreau, Alex G Ortega-Loayza

## Abstract

Pyoderma gangrenosum (PG) is a rare neutrophilic dermatosis causing chronic and recalcitrant painful ulcerations. Pathogenic mechanisms are yet poorly understood limiting therapeutic options, however, IL-12/IL-23 inhibition via ustekinumab has previously been associated with positive outcomes. We aimed to elucidate the dysregulated immune landscape of PG and lesional skin changes associated with IL-12/IL-23 blockade. We applied spatial transcriptomics and comparative computation analysis on lesional biopsies from two patients obtained before and after IL-12/IL-23 blockade with ustekinumab. Our data indicate lesional PG skin exhibits complex patterns of inflammation, including a not previously described major infiltration of B cells and establishment of tertiary lymphoid structures. In both patients, IL-12/IL-23 blockade led to marked clinical improvement but was associated with amelioration of contrasting inflammatory pathways. Notably, plasma cell markers and tertiary structures were recalcitrant to the treatment regime suggesting that B cells might play a role in the refractory nature of PG.

## Introduction

Pyoderma Gangrenosum (PG) is a rare and extremely painful neutrophilic dermatosis disease associated with non-healing cutaneous ulcers affecting 3-10 individuals per million people (1-5). The etiology and molecular pathogenetic events of this disease are yet unknown. Despite this unclear etiology, dysregulation of both innate and adaptive immunity has been implicated in PG, as supported by evidence of Th1/Th17 skewed inflammation and enhanced inflammasome activation associated with overexpression of neutrophils-derived cytokines (6-10). Appropriately targeting any of these associated pathways can regulate inflammatory milieu and may exhibit a therapeutic effect but recurrences are not uncommon (11-14). Dysregulation of IL (interleukin) 12 and IL-23 within the lesional dermis of PG has been implicated in PG pathogenesis (15-18) and here, we report two patients suffering from classic ulcerative PG and peristomal PG respectively, who eventually healed following IL-12/IL-23 inhibition via ustekinumab. To investigate how this therapeutic regime shaped the underlying inflammatory response, we performed spatial transcriptomics on lesional biopsies from patients before and after treatment. IL-12/IL-23 blockade correlated with reduction of some but not all underlying inflammatory pathways and revealed patterns of immune activation, including lymphocyte infiltration, formation of B cell rich tertiary lymphoid structures, and myeloid dysregulation.

## Results and Discussion

Two patients diagnosed with PG via PARACELSUS score refractory to TNF inhibition were treated with ustekinumab. Despite divergent clinical histories, both patents exhibited healing of their respective target ulcers (Figure 1A-D, Supplemental table 1). To investigate how the underlying immune response in these patients was shaped by ustekinumab, we performed spatial transcriptomics on lesional biopsies collected before and after treatment. Unsupervised clustering of pretreatment samples revealed both normal features of skin architecture (*i*.*e*., epidermal and dermal associated clusters) and a highly inflammatory milieu (Figure 2A, B). Notably, the patients exhibited broadly similar patterns of identified clusters with some specific differences in inflammatory regions (Figure 2B). Specifically, we identified four inflammatory clusters which were absent from previously published spatial transcriptomic annotations of normal human skin (Figure 2B) (19). All four clusters mapped to both patients but the relative abundance of each differed (Figure 2C).

**Figure 1.**
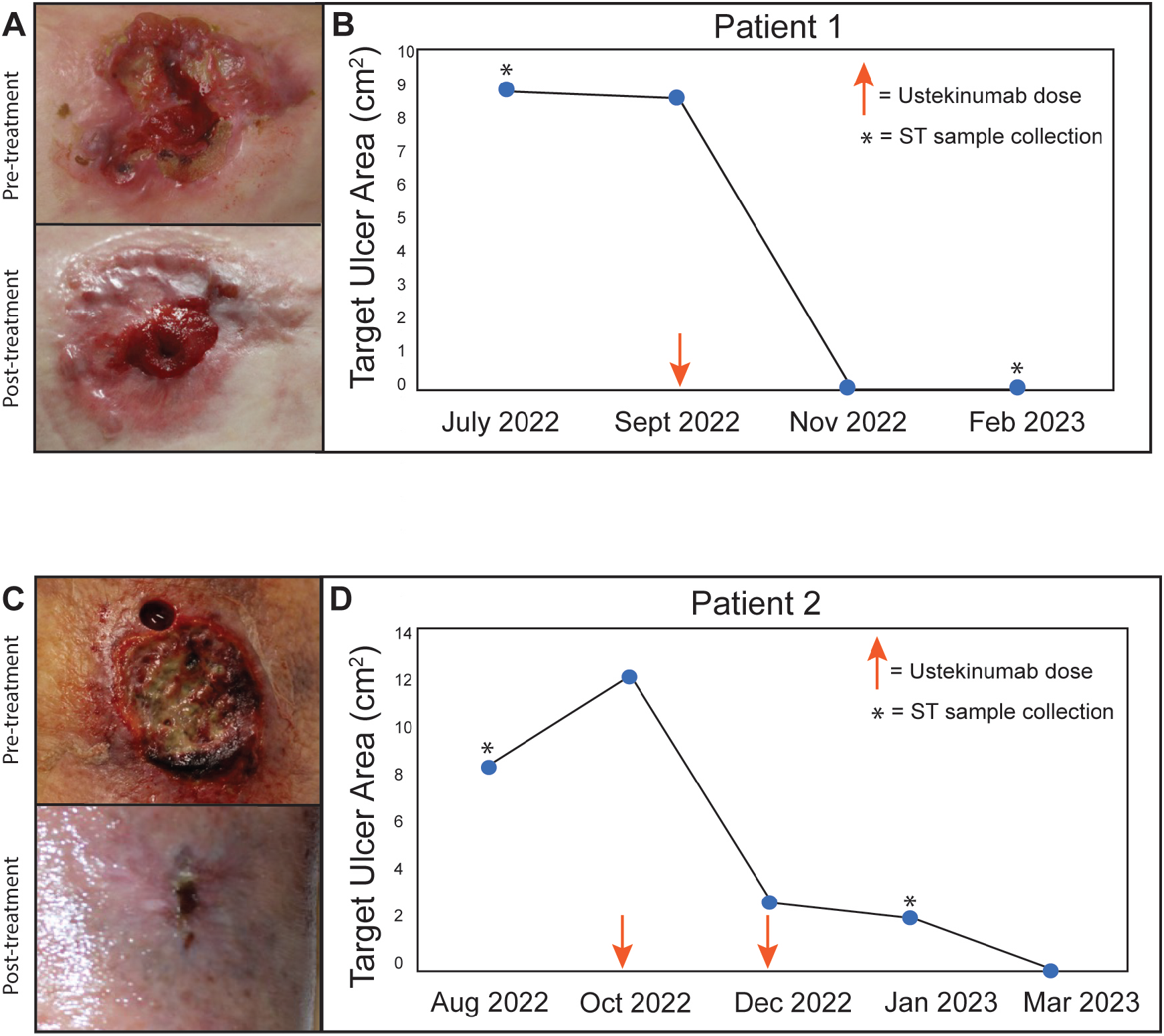
Target ulcer healing in pyoderma gangrenosum patients following ustekinumab treatment. **A)** Image of Patient 1 peristomal target ulcer before and after 90mg dose of ustekinumab treatment. **B)** Corresponding change in target ulcer area measured at every visit. **C)** Image of Patient 2 left lower extremity target ulcer pre- and post-treatment 90mg dose of ustekinumab treatment. **D)** Corresponding change in target ulcer area. Orange arrows correspond to ustekinumab doses. Asterisks denote the time of sample collection for ST.

**Figure 2.**
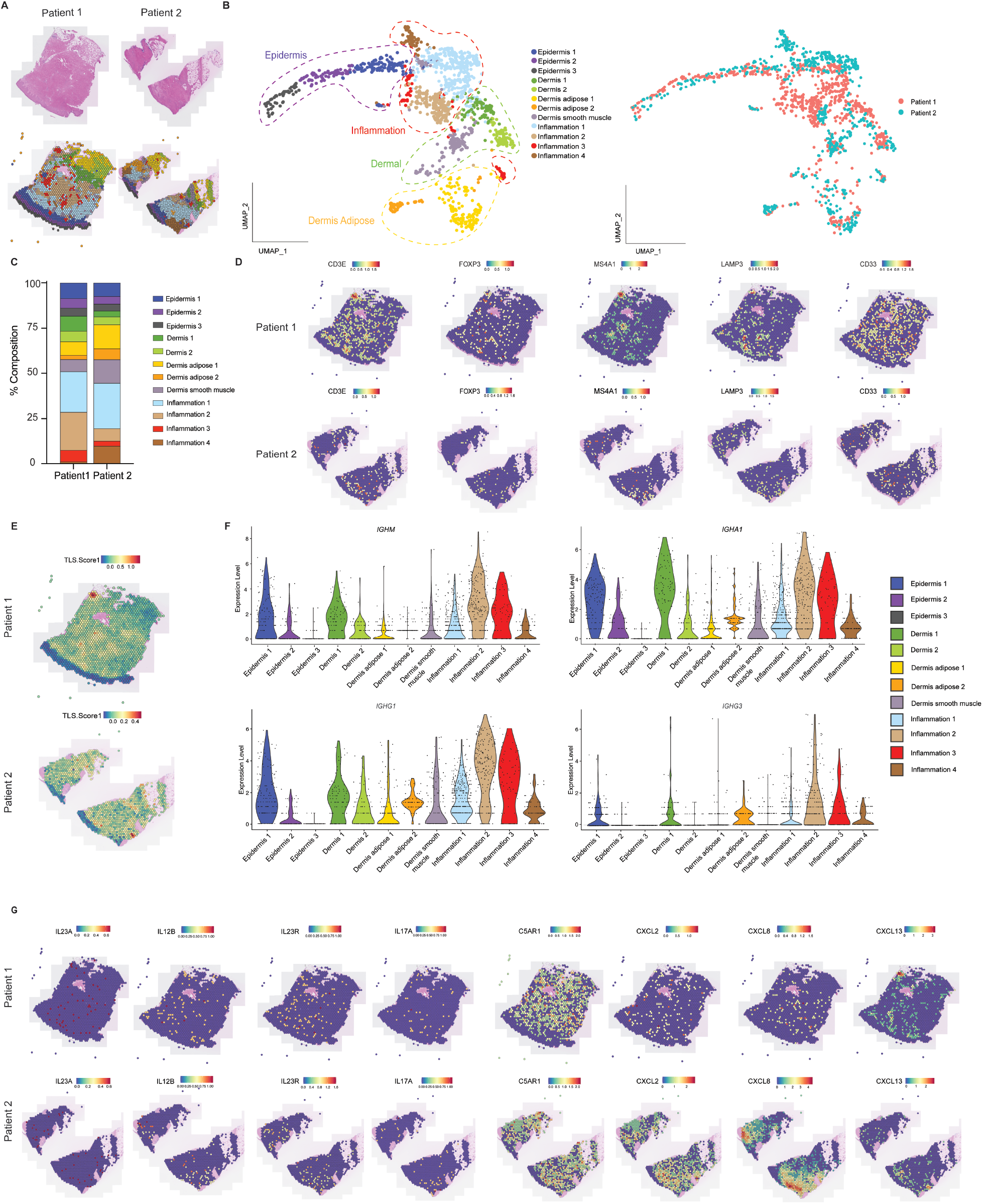
Spatial transcriptomic analysis of pyoderma gangrenosum lesions prior to ustekinumab treatment. **A**) Representative hematoxylin-eosin (H&E) (top) and corresponding unsupervised clustering spatial feature plot (bottom) following Harmony batch correction of both the patients. **B**) UMAP projection of 2535 spots colored by cluster obtained from two PG skin lesion combined (left) and separated by patients (right). **C**) Frequency plots exhibiting relative abundance of each cluster in both patients. **D**) Spatial feature plots of select leukocyte lineage genes. **E**) Spatial feature plot of TLS score in patient 1 (top) and patient 2 (bottom) derived from the top 50 TLS associated genes (20). **F**) Violin plots of immunoglobin genes expression per cluster of harmonized data from both patients. **G**) Spatial feature plots of select cytokine and chemokine genes.

Consistent with prior reports, tissue sections from both patients exhibited high levels of immune infiltrate and canonical lineage genes marked with the presence of T cells (*CD3E*), regulatory T cells (*FOXP3*), B cells (*MS4A1*), dendritic cells (*LAMP3*), and granulocytes (*CD33*) (Figure 2D) (12). In both patients, the spatial patterning of immune cells strongly coincided with the location of a tertiary lymphoid structure (TLS) gene signature (Figure 2E) (20). We further explored the enrichment of genes associated with class switched B cells and plasma cells and found high levels of *IGHM, IGHA1, IGHG1*, and *IGHG3* especially enriched in inflammation cluster 2 and 3 (Figure 2F). Analysis of cytokine and chemokines previously associated with PG pathogenesis revealed both conserved and discordant pathways between the two patients (21). Both patients exhibited high levels of *CS5AR1*, and the B cell chemoattractant *CXCL13*, while the second patient preferentially expressed *CXCL2* and *CXLC8* (Figure 2G). Thus, spatial transcriptomics of PG pretreatment samples revealed the presence of a diverse immune infiltrate, established TLS, plasma cell activity, and a complex network of inflammatory cytokines.

Spatial transcriptomic analysis of posttreatment lesional biopsies identified the same twelve clusters identified in baseline samples. This included clusters mapping to normal skin features, including epidermis and dermis, as well as inflammatory pathways (Figure 3A). However, a dynamic change in clusters was observed post-treatment in both patients (Figure 3B). We observed a striking decrease of inflammation cluster 1 in both patients, while inflammation 3 and 4 clusters diminished only in patient 1 and patient 2, respectively (Figure 3B). Pathway enrichment analysis revealed the dominant pathways associated with each cluster. Inflammation cluster 1 was linked with chemotaxis and innate immune activity, while inflammation clusters 2 and 3 were strongly associated with regulation of T and B cell activation, proliferation, and differentiation. Surprisingly, inflammation cluster 4 was largely driven by pathways implicated in sensory perception, including olfactory receptor genes, and hormone regulation (Figure 3C) In order to investigate how ustekinumab treatment regulated implicated cytokine and chemokine genes, we compared the 12 clusters between pre- and post-treatment samples in both patients. Pseudobulk differential expression analysis showed a compensatory increase in the expression of *IL23A, IL23R* and *IL-17A* post-treatment while the expression of *CXCL8* was significantly decreased suggesting this chemokine as a potential downstream target of IL-12/IL-23 blockade. Notably, expression of *TNFSF13B* (BAFF-a B cell survival factor) was enhanced post treatment. No changes in expression of *C5AR1, CXCL13* and *CXCL2* were observed (Figure 3D). Consistent with elevated *TNFSF13B*, TLS signatures remained strongly detectable in both the patients (Figure 3E). We further explored the treatment associated changes in B cells genes linked to immature B cells, class switched B cells, and plasma cells. While there was a reduction in *IGHM* genes post treatment, no such change was observed in *IGHA1, IGHG1* and *IGHG3* genes indicating that fully differentiated memory and plasma cells were retained in the lesional tissue (Figure 2F).

**Figure 3.**
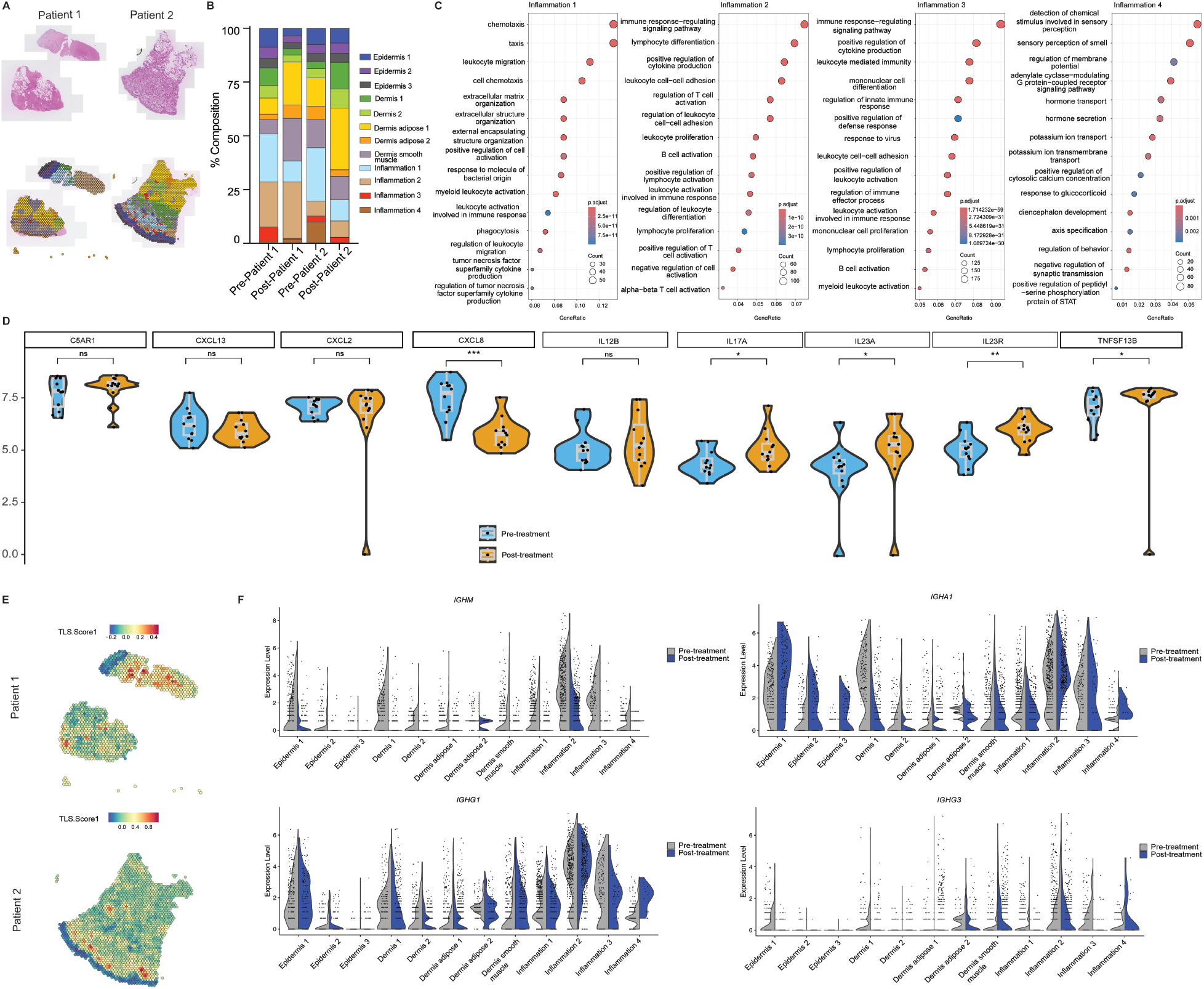
Spatial transcriptomic analysis of pyoderma gangrenosum lesions post-treatment. **A)** Representative hematoxylin-eosin (H&E) (top) and corresponding unsupervised clustering of ST plot (bottom) from 3142 spots following Harmony batch correction of both the patient’s post-treatment lesion. **B)** Composition plots exhibiting the relative abundance of each cluster of pre and post treatment of patient 1 and patient 2 utilizing a combined 5677 spots. **C)** Pathway enrichment analysis of differentially expressed genes in inflammation clusters. **D)** Pseudobulk analysis of genes in all 12 clusters from both patients comparing pre and post treatment groups **E**) Spatial feature plot of TLS addmodule score in patient 1 (top) and patient 2 (bottom) derived from the top 50 TLS associated genes (20). **F**) Violin plots of immunoglobin genes expression per cluster comparing pre and post treatment.

Although focused on only two patients, our study provides high resolution temporal insights into the inflammatory environment of PG lesional skin following treatment with IL-12/IL-23 blockade. This work highlights that while inhibition of representative cytokines of the Th1 and Th17 pathways shows promise as an additional alternative for some patients, it does not dampen all active inflammatory pathways. In particular, our data provides strong evidence that local B cell activation via formation of lesional TLS is recalcitrant to inhibition of the IL-12/IL-23 pathway. The persistence of class switched B cell implies that these cells may have a role in the refractory nature of some patients with PG and suggests another potential target for challenging patients.

## Methods

### Sample acquisition

Both patients were treated at the Department of Dermatology at Oregon Health and Science University (OHSU) and provided consent for tissue collection and wound photography under Oregon Health and Science IRB approved protocol #16753. Size 6mm punch biopsies from lesional and non-lesional tissue were obtained from both patients for analysis before and after ustekinumab treatment.

### Spatial Transcriptomics

Lesional skin punch biopsies were formalin-fixed and paraffin-embedded. Five-micron sections were placed on glass slides prior to hematoxylin-eosin staining and imaging with a Zeiss Axioscan 7 slider scanner. Spatial transcriptomics was performed with the Visium V2 Spatial Gene Expression kit (10X Genomics). Illumina sequencing was performed on prepared libraries by the OHSU Massively Parallel Sequencing Shared Resource using a NovaSeq 6000 (Illumina).

### Computational Analysis

SpaceRanger (v. 3.0) was used to process the FASTQ files and H&E-stained images using the Visium Human Transcriptome Probe Set v1.0 (GRCh38) followed by analysis with the Seurat R package (22). The files were normalized with SCTransform followed by spatial feature plot function, UMAP clustering and visualization (23). Samples were combined into a single, batch-corrected Seurat object prior to clustering using the R package Harmony (v.1.2.0) (24). The harmonized object was normalized and then dimensionality reduction, clustering, and visualization were performed. The FindAllMarkers function was used to generate a list of differentially expressed genes in each cluster. Only genes with a Log2FC greater than 0.25 and p-value less than 0.05 were kept. Clusters were manually annotated by comparing differentially expressed genes within spatial transcriptomics data generated by Naik et al, 2023 (19). We utilized their publicly available Seurat object “HEALTHY_ST_MARKERS.rds” and matched published annotations to analogous clusters in our harmonized object. TLS scores were calculated using the AddModuleScore function using gene sets described for TLSs (20). Enrichment analysis was derived using the package ‘clusterprofiler’ in R studio, using the function enrichGO (gene ontology) (25, 26). Differential genes were derived from the FindAllMarkers function in Seurat with arguments of only.pos = T, min.pct = 0.25, logfc.threshold = 0.25, test.use = ‘wilcox’. Pseudobulk analysis of genes “*IL23A*”, “*IL12B*”, “*IL23R*”, “*IL17A*”, “*C5AR1*”, “*CXCL2*”, “*CXCL8*”, “*CXCL13*”, “*TNFSF13B”* was generated using the AggregateExpression() function in the Seurat package in R Studio (23). Samples were grouped by treatment status and cluster for statistical analysis (t-test). Violin plots were generated using the ‘dittoSeq’ package in R studio (27).

